# AnnSQL: A Python SQL-based package for fast large-scale single-cell genomics analysis using minimal computational resources

**DOI:** 10.1101/2024.11.02.621676

**Authors:** Kenny Pavan, Arpiar Saunders

## Abstract

As single-cell genomics technologies continue to accelerate biological discovery, software tools that use elegant syntax and minimal computational resources to analyze atlas-scale datasets are increasingly needed. Here we introduce AnnSQL, a Python package that constructs an AnnData-inspired database using the in-process DuckDb engine, enabling orders-of-magnitude performance enhancements for parsing single-cell genomics datasets with the ease of SQL. We highlight AnnSQL functionality and demonstrate transformative runtime improvements by comparing AnnData or AnnSQL operations on a 4.4 million cell single-nucleus RNA-seq dataset: AnnSQL-based operations were executed in minutes on a laptop for which equivalent AnnData operations largely failed (or were ∼700x slower) on a high-performance computing cluster. AnnSQL lowers computational barriers for large-scale single-cell/nucleus RNA-seq analysis on a personal computer, while demonstrating a promising computational infrastructure extendable for complete single-cell workflows across various genome-wide measurements.

**Availability and Implementation:** AnnSQL is a pip installable package that can be found at https://github.com/ArpiarSaundersLab/annsql along with documentation at https://docs.annsql.com.

## 1. Introduction

Single-cell genomics technologies have emerged as transformative tools for discovering genome-wide RNA and chromatin landscapes of complex tissue at single-cell resolution in health and disease (Heumos et al., 2023; Nayak & Hasija, 2021; Wen et al., 2022). Popular software tools have converged on data structures that facilitate storage, preprocessing and myriad downstream analyses by prioritizing organizational clarity at the cost of computational performance. For example, Scanpy (Wolf et al., 2018) and the Scverse ecosystem (Virshup et al., 2021) use AnnData, while Seurat (Butler et al., 2018) uses Seurat objects. Now routine, “atlas-scale” descriptions of millions of cells based on tens of thousands to millions of genomics features strain software tools that use hierarchical structures (like AnnData) and often require high-performance computing clusters to execute. We were inspired to create AnnSQL for simple analysis of large datasets because in other contexts, SQL databases are a popular storage choice due to high transactional speed of row-oriented storage formats (Plattner, 2009) and because no other SQL-based single-cell genomics analysis packages currently exist.

H5 hierarchical objects like AnnData have become standard storage for single-cell genomics software development in recent years (The HDF Group, 2020/2024). AnnData provides a convenient and compact structure for storing, organizing and analyzing high-dimensional datasets. Applications can load AnnData entirely into system memory or, for large datasets, stream AnnData from disk using the “backed mode” parameter. Although AnnData allows faster statistical operations than traditional row-oriented databases, the backed mode often lacks support for aggregate functions.

Advances in SQL databases present new opportunities for single-cell genomics applications. Traditional SQL databases are limited because they often require technical knowledge for configuring and running databases as a background system process that requires connection to the service via the application, adding complexity that restricts the ease of data exploration. As an alternative, SQLite (Gaffney et al., 2022) operates in-process by creating a file-based database capable of direct application access yet a reliance on row-oriented transactions can restrict performance on statistical operations. To enable high-performance statistical functions and in-process queries, DuckDb (Raasveldt & Mühleisen, 2019) was implemented as a column-based storage engine that operates using chunked vectors of column data. Strict in-process operation allows DuckDb data to be stored in a file and accessed in a memory-efficient manner using SQL via the Python API. These two database engines have recently been profiled for use with several genomic file formats, without exploring scRNA-seq data (Kioroglou, 2025). We believe the features of DuckDb nominate it as a powerful storage and processing engine for cell by gene feature matrices.

We developed AnnSQL to bridge the gap between AnnData objects and SQL databases for single-cell genomics analysis. AnnSQL converts each layer of the AnnData hierarchical object into an equivalent SQL table, using the Python DuckDb API. AnnSQL supports both in-memory and backed modes of AnnData, enabling larger than memory databases to be built with minimal computational resources. AnnSQL-instantiated datasets can be fluidly queried using SQL syntax, allowing aggregate and statistical functions to be performed on larger than memory datasets with exceptionally fast run times.

## 2. Results

### 2.1 Package Capabilities

The AnnSQL package enables SQL-based queries on AnnData objects, returning results as either an AnnData object, Pandas dataframe (McKinney, 2010), or Parquet file (Vohra, 2016), thus seamlessly supporting a variety of downstream analysis tools or pipelines. Functionally, AnnSQL captures the AnnData hierarchical H5 data structure and represents the data in a relational database using the in-process DuckDb engine (**Figure 1A**). AnnSQL supports in-memory and on-disk database representations, useful for smaller or larger datasets, respectively. AnnSQL databases are built using the *MakeDb* class that supports AnnData inputs from both backed and non-backed modes. Once a database has been created and instantiated using the AnnSQL class (**Figure 1B, 1C**), the AnnSQL package provides methods for querying, updating, and deleting cell or genome-feature data, as well as extended functionality to perform common single-cell manipulations or analyses, such as normalization, log transforms, determining library or gene counts, calculating feature variance, principal component analysis, and differential expression.

**Figure 1.**
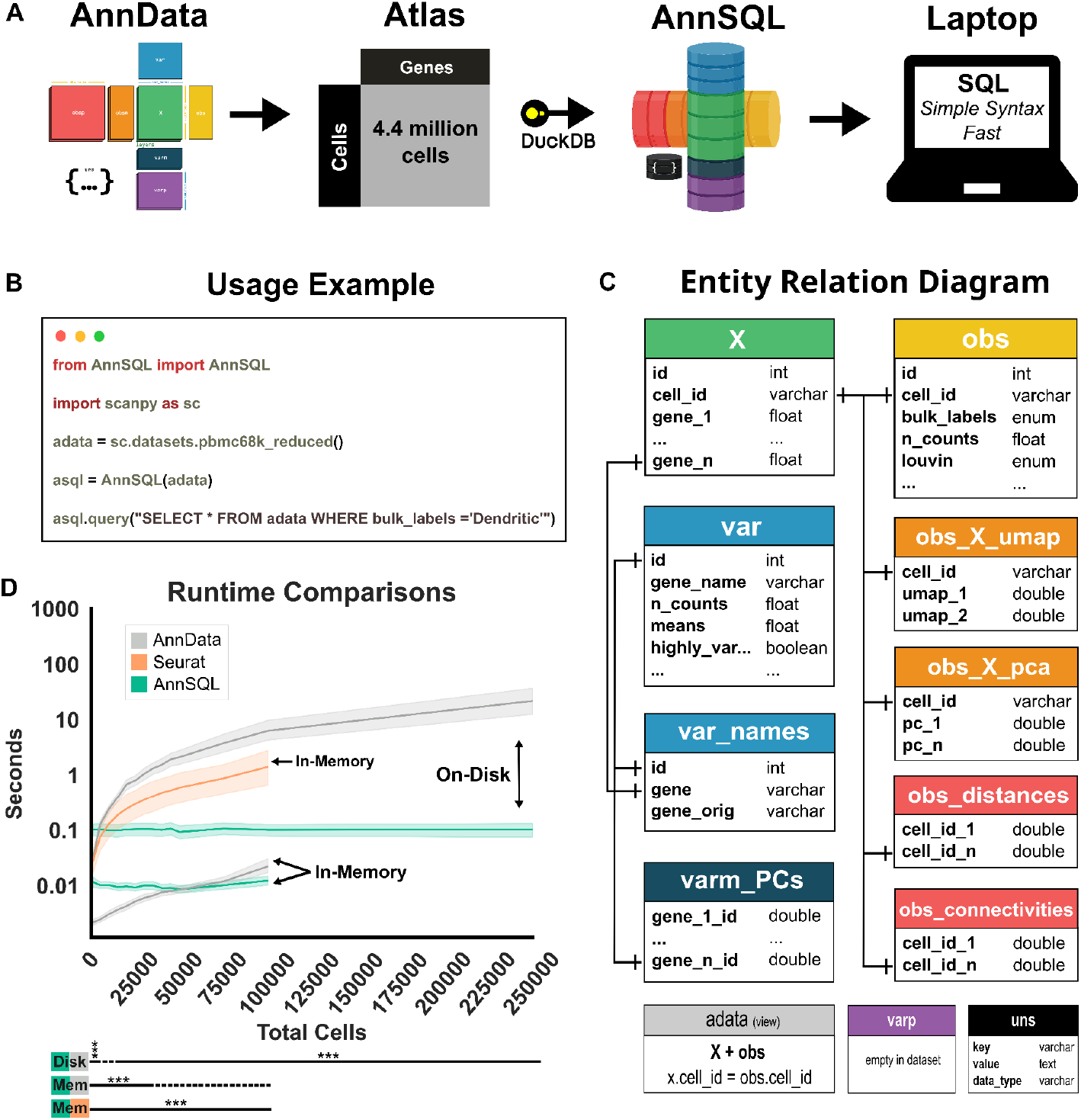
A) AnnSQL workflow. AnnSQL converts AnnData objects using DuckDB to enable high-performance SQL queries. B) Basic usage example using the pbmc68k reduced dataset provided with Scanpy. C) The ERD of the database generated from the pbmc68k reduced dataset. D) Runtime comparisons of six queries and filters (**Figure S1A**) for AnnSQL,AnnData (backed and non-backed modes displayed as on-disk and in-memory, respectively), and Seurat objects (in-memory) repeated six times for each of the six queries and filters at each library size. Shading represents 95% confidence intervals. Bottom lines represent statistical significance of each library size with respect to the comparison. Solid lines indicate significant, dashed lines indicate not significant (***p < 0.001; paired t-test; Bonferroni corrected).

### 2.2 Runtime Estimates

To systematically compare runtime differences across AnnData filters and AnnSQL queries, we generated six increasing complex queries of synthetic scRNA-seq data and stratified our analysis by in-memory or on-disk modes for each method (**Figure 1D** and **S1A**)(**Methods**). In-memory comparisons trials ranged from 1,000 to 100,000 cells before reaching system memory limits, while on-disk trials ranged from 1,000 to 250,000 cells. Each dataset contains a fully dense matrix (**Methods**). Critically, AnnSQL runtimes were essentially unchanged across the full range library sizes tested for both in-memory and on-disk. On-disk AnnData outperformed on-disk AnnSQL only for libraries < 5,000 cells, while queries of 250,000 cells were on average ∼400 times faster using AnnSQL (0.14 versus 57 seconds). In-memory AnnData outperformed in-memory AnnSQL only for libraries with < 45,000 cells and both methods achieved <0.01 second runtimes with <75,000 cell libraries. Additionally, we repeated the same experiment using *in silico* datasets generated by the Splatter (Zappia et al., 2017) sc/nRNA–seq simulation tool. Runtimes of all comparisons decreased as each of these datasets contain sparser UMI counts (**Supplemental 2A**). These comparative runtime metrics illustrate the drastic speed improvements and size-scalability of AnnSQL processing of large scRNA-seq datasets.

### 2.3 Use Case: Querying and processing 4.4 million cells on a laptop

To illustrate AnnSQL runtime improvements, we analyzed a single nucleus RNA-seq atlas of the mouse containing 4.4 million cells and annotations (Langlieb et al., 2023). First, we opened the atlas AnnData object in backed mode and created an on-disk AnnSQL database using the *MakeDb* class (**Methods**). Next, we performed routine procedures (cell filtering, summing library feature counts, cell-count normalization and cell-count log transformation) using either the original AnnData object (in-memory or on-disk/backed mode) or on-disk AnnSQL (using methods provided in our package) on both a laptop and a High Performance Cluster (HPC) (**Table 1** and **Methods**). Building a Seurat object based on the atlas dataset failed due to memory errors and for all procedures other than filtering, AnnData analysis failed on both the laptop or HPC due to a lack of support for backed-mode or memory errors. Equivalent procedures using AnnSQL completed between ∼4.28-67 minutes on a laptop and were up to 3x faster than on the HPC. While backed-mode (but not in-memory) filtering could be accomplished on a laptop using AnnData, this procedure was ∼746x faster using AnnSQL. The HPC did enable filtering using in-memory AnnData, but was still ∼42x faster on a laptop with AnnSQL. These data illustrate how AnnSQL performance enhancements allow users to access and manipulate atlas-scale scRNA-seq data from their personal computer with ease.

**Table 1:** Runtime and memory usage (mean ± SD; n=5) of procedures for equivalent AnnSQL and AnnData objects containing 4.4 million cells. Converting the dataset to a Seurat object failed due to memory constraints. Scanpy preprocessing functions were used for AnnData, while AnnSQL extended functionality methods were used for the AnnSQL database. Memory Error indicates the run exceeded available memory; Not Supported indicates the procedure is not compatible with AnnData in backed mode.

| Procedure | Approach | Laptop |  | HPC (sec) |  |
| --- | --- | --- | --- | --- | --- |
|  |  | Runtime (sec) | Memory (Mb) | Runtime (sec) | Memory (Mb) |
| Filtering | AnnSQL | 0.23 ±0.01 | 147.29 ±34.2 | 0.99 ±0.14 | 321.64 ±2.76 |
|  | AnnData (Backed) | 171.62 ±11.3 | 89.6 ±0.13 | 149.03 ±17.47 | 788.51 ±67.32 |
|  | AnnData (Memory) | Memory Error |  | 9.59 ±3.13 | 368433.14 ±165.17 |
|  | Seurat | Memory Error |  | Memory Error |  |
| Total Library Counts | AnnSQL | 257.35 ±8.04 | 3799.4 ±10.11 | 446.73 ±8.28 | 4347.44 ±52.62 |
|  | AnnData (Backed) | Not Supported |  | Not Supported |  |
|  | AnnData (Memory) | Memory Error |  | Memory Error |  |
|  | Seurat | Memory Error |  | Memory Error |  |
| Normalize Expression | AnnSQL | 3924.7 ±59.27 | 29407.81 ±21.51 | 10922.57 ±178.22 | 56965.75 ±29.98 |
|  | AnnData (Backed) | Not Supported |  | Not Supported |  |
|  | AnnData (Memory) | Memory Error |  | Memory Error |  |
|  | Seurat | Memory Error |  | Memory Error |  |
| Log Transform | AnnSQL | 4057.37 ±65.0 | 29879.29 ±21.7 | 12739.63 ±166.97 | 29813.2 ±124.56 |
|  | AnnData (Backed) | Not Supported |  | Not Supported |  |
|  | AnnData (Memory) | Memory Error |  | Memory Error |  |
|  | Seurat | Memory Error |  | Memory Error |  |

## 3. Methods

### 3.1 Package Software Structure

AnnSQL was built as an object-oriented Python package that parses the layer structure of typical AnnData objects into SQL tables (**Figure 1C**) and provides flexible parameters. All functionality presented in AnnSQL v1, exists in 3 classes: (1) The *BuildDb* class is used to construct both in-memory and on-disk databases using the DuckDb Python client API. (2) The *MakeDb* class can be instantiated to create an on-disk database from a properly formatted AnnData object and stored with the “.asql” file extension. (3) The class *AnnSQL* is the main access point to instantiate and interrogate a dataset. The package has methods for queries, allows multiple return types, and contains extended functionality to help with data preprocessing.

### 3.2 Database Construction

Each layer in a common single-cell AnnData object is a parameter in the AnnSQL and *MakeDb* classes. Both classes make calls to the *BuildDb* class to construct a relational database with the AnnData layers determined by the user (*X, obs, var, var_names, obsm, varm, obsp*, and *uns). X* is created as a table where each gene is a column cast as a FLOAT with the addition of a “cell_id” column to contain the cell barcode. The *obs* table structure is composed of columns for the “cell_id” and each *obs* property. The *var* table contains the vertical layer associated with the AnnData object where the auto incremented id represents the gene and each column represents *var* layer properties. Each *obsm, varm*, and *obsp* layer property is composed of a different data type or structure. To convert layers to SQL tables, we created a new table for each layer property. For example, the X_pca property of the *obsm* layer is translated into the table, “obsm_X_pca” containing the cell by PCA matrix. Lastly, the *uns* AnnData layer is serialized and inserted into the “*uns_raw*” table. This table contains the property name defined by the key column, a serialized data value column, and the original data_type column, such that the data can be reconstructed. Lastly, a convenience view “*adata*” is created by joining the obs and X table. Developers can optionally bypass the view creation by using the “*convenience_view*” boolean parameter. Lastly, special characters in gene names or other table columns are stripped of non-sql safe characters and replaced with underscores.

### 3.3 Database Query Wrapper

The AnnSQL class contains several methods for updates, deletes, and queries. The class requires either a path to a previously built AnnSQL database or an AnnData object to load into memory. After instantiation, the *query* method interfaces with the database to return a pandas dataframe (default), AnnData, or Parquet file. The *query_raw* method opens a db connection and executes a SQL statement. Additionally, the methods *show_settings* and *show_tables* exist for convenience.

### 3.4 Extended Functionality

AnnSQL contains functionality for basic preprocessing as methods within the AnnSQL class which include: Total counts per library, counts per gene, data normalization, log transformations, and gene expression variance calculation, PCA calculation, UMAP, Leiden clustering, and differential expression; along with a variety of helper methods and plotting utilities; all documented at https://docs.annsql.com. To accomplish data total counts per library, ***calculate_total_counts*** iterates cells in chunks and adds the “total_counts” column to the *X*, then *obs* table. Calculating total gene counts is accomplished in the ***calculate_gene_counts*** method by adding a “total_gene_counts” column to the *var* table and using the internal SUM function. The ***expression_normalize*** method updates the *X* table in chunks by dividing gene counts by “total_counts” then multiplying by the normalized value (default: 10^4^). The ***expression_log*** method updates all values in the *X* table with the user defined ln, log2 or log10 parameter options. Lastly, ***calculate_variable_genes*** uses the population variance function in DuckDb to calculate gene-specific expression and stores the result in the “variance” column of the *var* table. In chunk parameters, gene sample variance is calculated including Bessel’s bias correction. The ***calculate_pca*** method determines principal components (PCs) for the data stored in the specified table. This method uses the top variable genes to perform PCA. The PCA calculation is performed as a hybrid of SQL and Python to create a covariance matrix, compute eigenvalues and eigenvectors which are used to determine PCs for each cell. The results are stored in the PC_scores table and can be queried. After reducing the data size using PCA, we opted to implement UMAP and Leiden clustering methods as wrapper methods to existing “umap-learn” and “scikit-network” pip packages, respectively. The results of these operations are stored in the umap_embeddings table and as the obs.leiden_clusters field. Detailed descriptions of additional functionality can be found at https://docs.annsql.com.

### 3.5 Runtime Analyses

Runtime tests were performed on simulated and Mouse Brain Atlas data (Langlieb et al., 2023). Simulated AnnData objects contained random expression values and saved as h5ad files ranging from 1,000 to 250,000 cells. Each file contained 10,000 genes and an *obs* layer with random cell type assignments. File queries and filters were compared (**Figure S1A)** and generated the runtime results (**Figure 1D)**. Additionally, a second set of *in silico* datasets were generated using Splatter, a sc/nRNA-seq simulator, with the same shapes as the randomly generated data. We applied the same analysis to these datasets and presented our findings in **Supplemental 2A**.

The Mouse Brain dataset was converted into an on-disk AnnSQL database using the *MakeDb* class, then queried and processed as described in **Table 1**. Filters are as follows:

### AnnData

*adata[adata[:, “ENSMUSG00000070880”]*.*X > 0, “ENSMUSG00000070880”]*

### AnnSQL

*asql*.*query(“SELECT ENSMUSG00000070880 FROM X WHERE ENSMUSG00000070880 > 0”)*

Runtime analyses were performed on an Ubuntu 24.04 laptop, containing 40Gb memory, 1Tb SSD Drive and a 12th Gen Intel® Core™ i7-1255U×12 CPU or a HPC single node containing 46 CPUS (Intel® Xeon® Gold 6542Y), 512Gb memory and high-performance storage. Runtime scripts are in the “examples/analyses” repository.

## 4. Discussion

Here, we introduce AnnSQL for analysis of single-cell genomics data using SQL syntax and showcase query and preprocessing performance using large datasets with minimal computational resources. AnnSQL uses the in-process DuckDb engine and provides extended functionality to showcase how vectorized queries of the column-based storage engine benefit single-cell genomics applications. In future releases, extended functionality methods can be further improved by utilizing DuckDb user-defined functions. Additionally, the DuckDb client API supports programming languages popular in genomics research and development (including Java, R, C++, Julia, and cmd libraries).

AnnSQL runtime comparisons highlight impressive performance improvements achieved that help democratize access to large-scale single-cell RNA-seq data, enabling laptop-based analysis of millions of cells datasets. The ease and clarity of SQL syntax, along with Python-based wrapper functions, further lowers the analysis barriers. We do note that for small-scale datasets (<45,000 cells), AnnData slightly outperforms AnnSQL operations, but minor differences in runtime (< 0.1 seconds) might be tolerated if users prefer SQL-based data access.

In developing AnnSQL, we intentionally mirrored the AnnData layered structure for interpretability. While an ideal H5 based structure, AnnData is a non-optimal structure for a relational database (Codd, 1970). Future work converting AnnData objects into an optimized normal formed database may further decrease runtimes (Kent, 1983).

While the use of SQL for single-cell genomics data storage and analysis has yet to be extensively explored, our results indicate that SQL-based databases may be increasingly valuable as datasets continue to grow. We suspect this lack of attention was due to either the technical knowledge necessary for database configuration or the resources necessary to process a cell by genomic feature matrix in a typical implementation (such as SQLite or MySql (*MySQL*, n.d.)). Our AnnSQL results suggest column-based SQL approaches for filtering or processing single-cell data should be considered when (1) fluid SQL syntax is preferred; (2) dataset size exceeds system memory; or (3) when minimal computational resources are desired for exploring large datasets without downsampling.

## Author Contributions

Conceptualization (KP); Data Curation (KP); Formal Analysis (KP); Funding acquisition (AS); Investigation (KP); Methodology (KP, AS); Project Administration (AS); Resources (KP, AS); Software (KP); Supervision (AS); Validation (KP); Visualization (KP); Writing – original draft (KP); Writing – review & editing (KP, AS).

## Acknowledgements

The authors thank Connor Frankston, Zach Goode, Dr. Guanming Wu, Dr. Lamya Ben Ameur in addition to the members of the Saunders lab, Andrew Adey lab, and Brian O’Roak lab for helpful advice and/or discussions, as well as Theresa Provitola for design assistance with the AnnSQL logo.

---

**Supplemental 1A.**
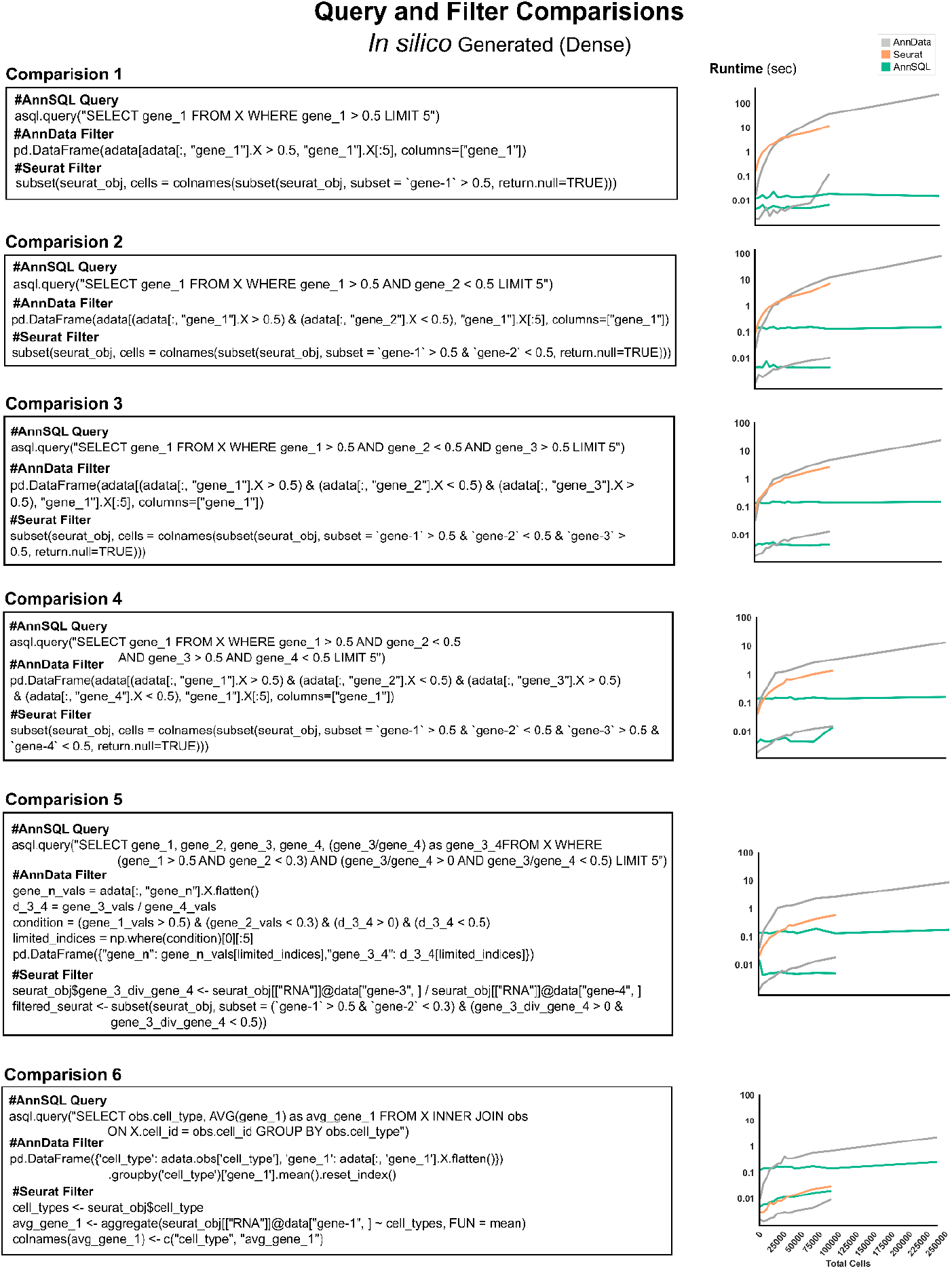
Runtime comparisons for six equivalent operations (rows) performed using AnnData versus AnnSQL (summarized in Figure 1D). Left, operation-specific calls. Right, runtime comparisons for on-disk and in-memory modes.

**Supplemental 2.**
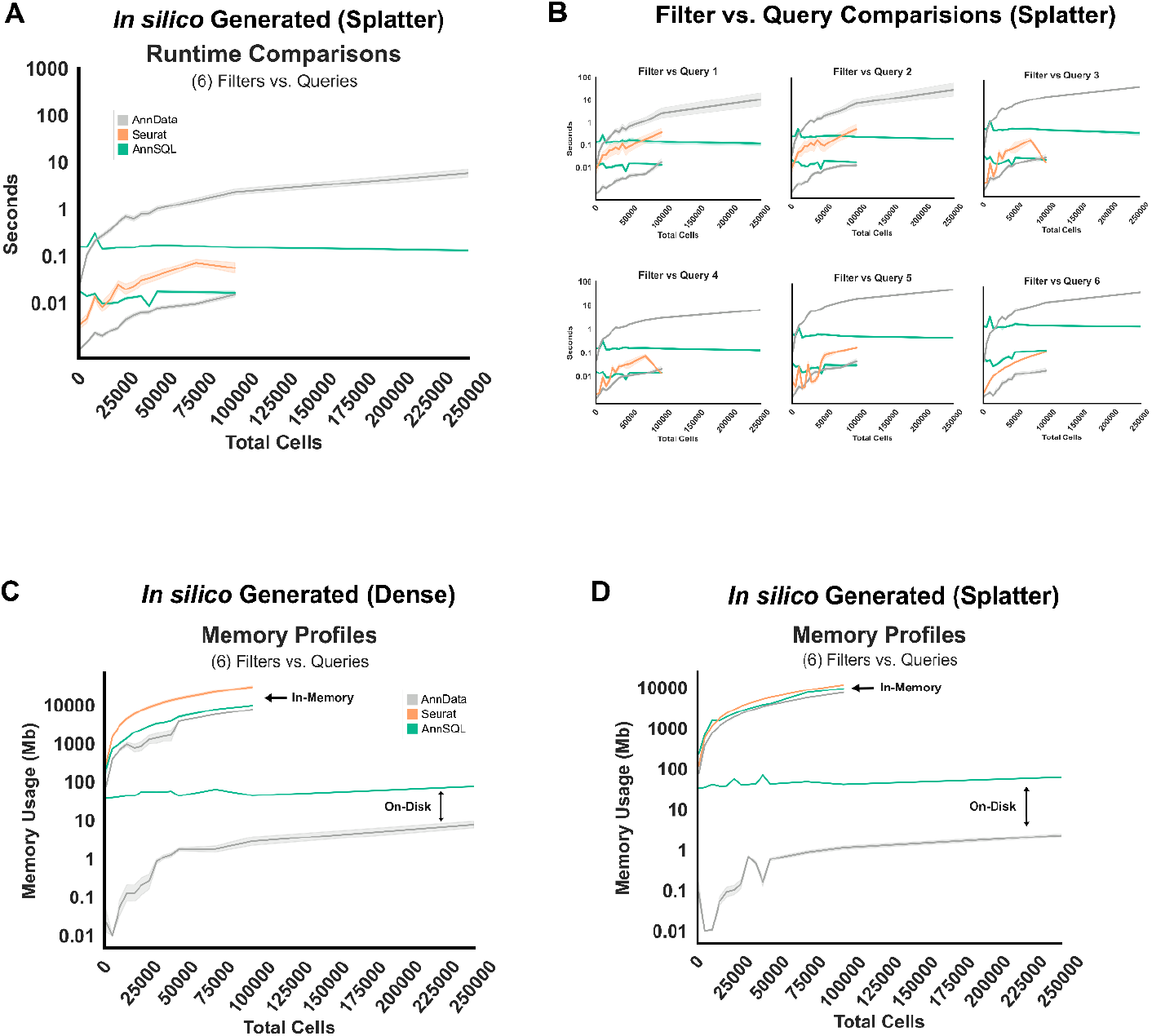
A.) *In silico* runtime analysis of libraries generated using the Splatter sc/nRNA-seq simulator tool. Filter and query comparisons are identical to **Supplemental 1A**. B.) Individual runtime profiles across library sizes for individual filters 1-6 of Splatter generated data. C.) Memory consumption of dense in-silico generated while running query and filter comparisons. D.) Same as C but with Splatter (sparse) generated data. For each figure, filters and queries are repeated six times for each of the six queries and filters. Shading represents 95% confidence intervals.

